# Delta-band Cortical Tracking of Acoustic and Linguistic Features in Natural Spoken Narratives

**DOI:** 10.1101/2020.07.31.231431

**Authors:** Cheng Luo, Nai Ding

**Affiliations:** Key Laboratory for Biomedical Engineering of Ministry of Education, College of Biomedical Engineering and Instrument Sciences, Zhejiang University, Hangzhou, China 310027; Research Center for Advanced Artificial Intelligence Theory, Zhejiang Lab, Hangzhou 311121, China

## Abstract

Speech contains rich acoustic and linguistic information. During speech comprehension, cortical activity tracks the acoustic envelope of speech. Recent studies also observe cortical tracking of higher-level linguistic units, such as words and phrases, using synthesized speech deprived of delta-band acoustic envelope. It remains unclear, however, how cortical activity jointly encodes the acoustic and linguistic information in natural speech. Here, we investigate the neural encoding of words and demonstrate that delta-band cortical activity tracks the rhythm of multi-syllabic words when naturally listening to narratives. Furthermore, by dissociating the word rhythm from acoustic envelope, we find cortical activity primarily tracks the word rhythm during speech comprehension. When listeners’ attention is diverted, however, neural tracking of words diminishes, and delta-band activity becomes phase locked to the acoustic envelope. These results suggest that large-scale cortical dynamics in the delta band are primarily coupled to the rhythm of linguistic units during natural speech comprehension.

## Introduction

When listening to speech, low-frequency cortical activity in the delta (< 4 Hz) and theta (4-8 Hz) bands is phase locked to speech (Keitel et al., 2018; Luo and Poeppel, 2007). However, it remains debated what speech features are encoded in the low-frequency cortical response. A large number of studies have demonstrated that the low-frequency cortical response is correlated with low-frequency power fluctuations in speech, i.e., the speech envelope (Destoky et al., 2019; Ding and Simon, 2012; Koskinen and Seppa, 2014; Liberto et al., 2015; Nourski et al., 2009; Peelle et al., 2013). Since the speech envelope provides an important acoustic cue for syllable boundaries, it has been hypothesized that neural activity tracking the speech envelope is a mechanism to segment continuous speech into discrete units of syllables (Giraud and Poeppel, 2012; Poeppel and Assaneo, 2020). In other words, the envelope tracking neural response reflects an intermediate neural representation linking the auditory representation of acoustic speech features and phonological representation of syllables. Consistent with this hypothesis, it has been found that neural tracking of speech envelope is both tied to low-level speech features (Doelling et al., 2014) and related to perception: On the one hand, it can occur when speech recognition fails (Etard and Reichenbach, 2019; Howard and Poeppel, 2010; Pena and Melloni, 2012; Zoefel and Vanrullen, 2016; Zou et al., 2019). On the other hand, it strongly modulated by attention (Golumbic et al., 2013; Kerlin et al., 2010) and may be a prerequisite for successful speech recognition (Vanthornhout et al., 2018).

Recently, evidence also shows that delta-band cortical activity can directly track multiple levels of linguistic units, e.g., words and phrases, even if these linguistic units have no acoustic correlates (Brodbeck et al., 2018; Broderick et al., 2018; Buiatti et al., 2009; Ding et al., 2016a; Jin et al., 2018; Makov et al., 2017; Sheng et al., 2019). Based on these phenomena, it has been hypothesized that delta-band cortical activity can reflect linguistic-level neural representations that are constructed based on internal linguistic knowledge instead of acoustic cues (Ding et al., 2016a; Ding et al., 2018). Nevertheless, to dissociate linguistic units with the related acoustic cues, the studies showing linguistic tracking responses mostly employ well-controlled synthesized speech that is presented as an isochronous sequence of syllables. Furthermore, these studies mostly present a sequence of unrelated linguistic units, the boundaries between which are perceptually more salient than the linguistic boundaries in natural speech. Therefore, it remains unclear whether neural activity can track linguistic units in natural speech, which is semantically coherent but not periodic, containing both acoustic and linguistic information in the delta band.

Here, we first asked whether cortical activity could track disyllabic words composed of two monosyllabic morphemes in semantically coherent stories. The story was either naturally read or synthesized as an isochronous sequence of syllables as in previous studies that demonstrate linguistic structure tracking (Ding et al., 2016a). We then asked whether neural tracking of disyllabic words interacted with neural tracking of delta-band acoustic envelope. To address this question, we introduced an artificial delta-band envelope to isochronous speech and tested how it influenced the word-tracking response. Finally, since previous studies have shown that cortical tracking of speech strongly depended on the listeners’ task (Ding and Simon, 2012; Golumbic et al., 2013; O’Sullivan et al., 2015), we recorded the neural responses during both active speech comprehension and unattended listening.

## Results

### Neural tracking of words in isochronously presented narratives

We first presented semantically coherence stories that were synthesized as an isochronous sequence of syllables (Fig. 1A, left). For the stories with a metrical structure, every other syllable must be a word onset. More specifically, the odd terms in the metrical syllable sequence were always the initial syllable of a word, while the even terms were either the second syllable of a disyllabic word (73% probability) or a monosyllabic word (23% probability). In the following, the odd terms of the syllable sequence were referred to as σ1, while the even terms were referred to as σ2. Since the syllables were presented at a constant rate of 4 Hz, the neural response tracking syllables was frequency tagged at 4 Hz. Furthermore, since every other syllable in the sequence was the onset of a word, neural activity phase locked to word onsets was expected to show a regular rhythm at half of the syllabic rate, i.e., 2 Hz (Fig. 1A, right).

**Fig. 1.**
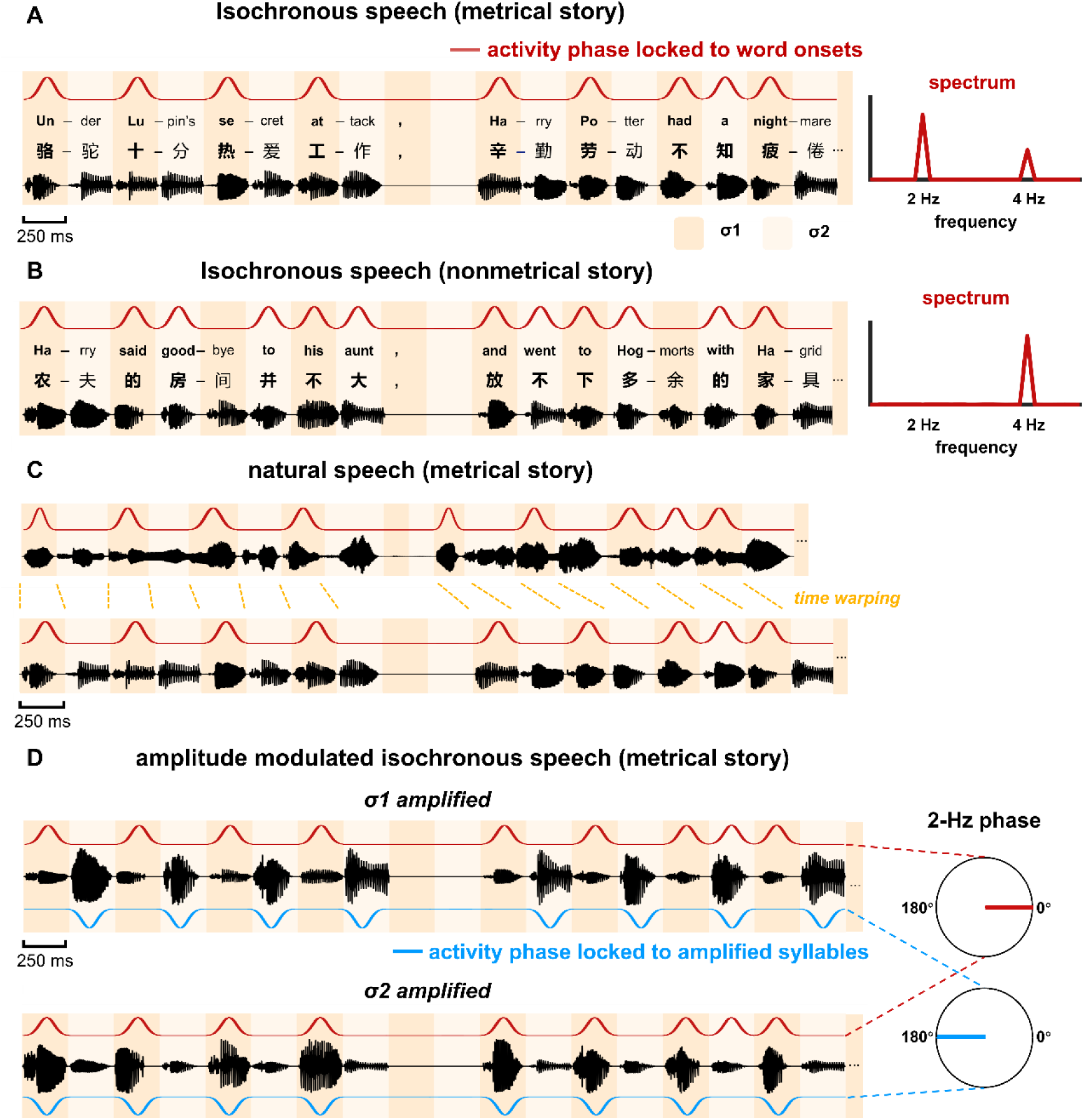
Stimulus. (AB) Two types of stories are constructed: metrical stories and nonmetrical stories. (A) Metrical stories are composed of disyllabic words and pairs of monosyllabic words, so that the odd terms in the syllable sequence (referred to as σ1) must be the onset of a word. Here the onset syllable of each word is shown in bold. All syllables are presented at a constant rate of 4 Hz. A 500-ms gap is inserted at the position of any punctuation. The red curve illustrates cortical activity that is phase locked to word onsets. It shows a 2-Hz rhythm, which can be clearly observed in the spectrum shown on the right. The stories are in Chinese and the English examples are shown for illustrative purposes. (B) In the nonmetrical stories, word onsets are not regularly positioned, and activity that is phase locked to word onsets does not show 2-Hz rhythmicity. (C) Natural speech. The stories are naturally read by a human speaker and the duration of syllables is not constrained. During the analysis, the response to natural speech is time warped to simulate the response to isochronous speech, and the time-warped word-tracking response is expected to show a 2-Hz rhythm. (D) Amplitude modulated isochronous speech is constructed by amplifying either σ1 or σ2 by a factor of 4, creating a 2-Hz acoustic envelope. The red and blue curves illustrate responses that are phase locked to word onsets and amplified syllable, respectively. The word-tracking response is identical for σ1- and σ2-amplified speech, while the response tracking the amplified syllable shows a 180° degree phase shift between conditions.

As a control condition, we also presented stories with a nonmetrical structure (Fig. 1B). These stories were referred to as the nonmetrical stories in the following. In these stories, the word duration was not constrained and σ1 was not always a word onset. Consequently, the word onsets in these stories did not show rhythmicity at 2 Hz, and neural activity phase locked to word onsets was not frequency tagged at 2 Hz.

All stories were presented to two groups of participants who performed different tasks. One group of participants was asked to attend to the stories and answer 3 comprehension questions after each story. The participants correctly answered 96% ± 9% and 94% ± 9% questions for metrical and nonmetrical stories, respectively. The other group of participants, however, was asked to watch a silent movie while listening and did not have to answer any speech comprehension question. The silent movie was not related to the auditorily presented stories. The EEG responses to isochronously presented stories were shown in Fig. 2AB. The response spectrum was averaged over participants and EEG channels.

**Fig. 2.**
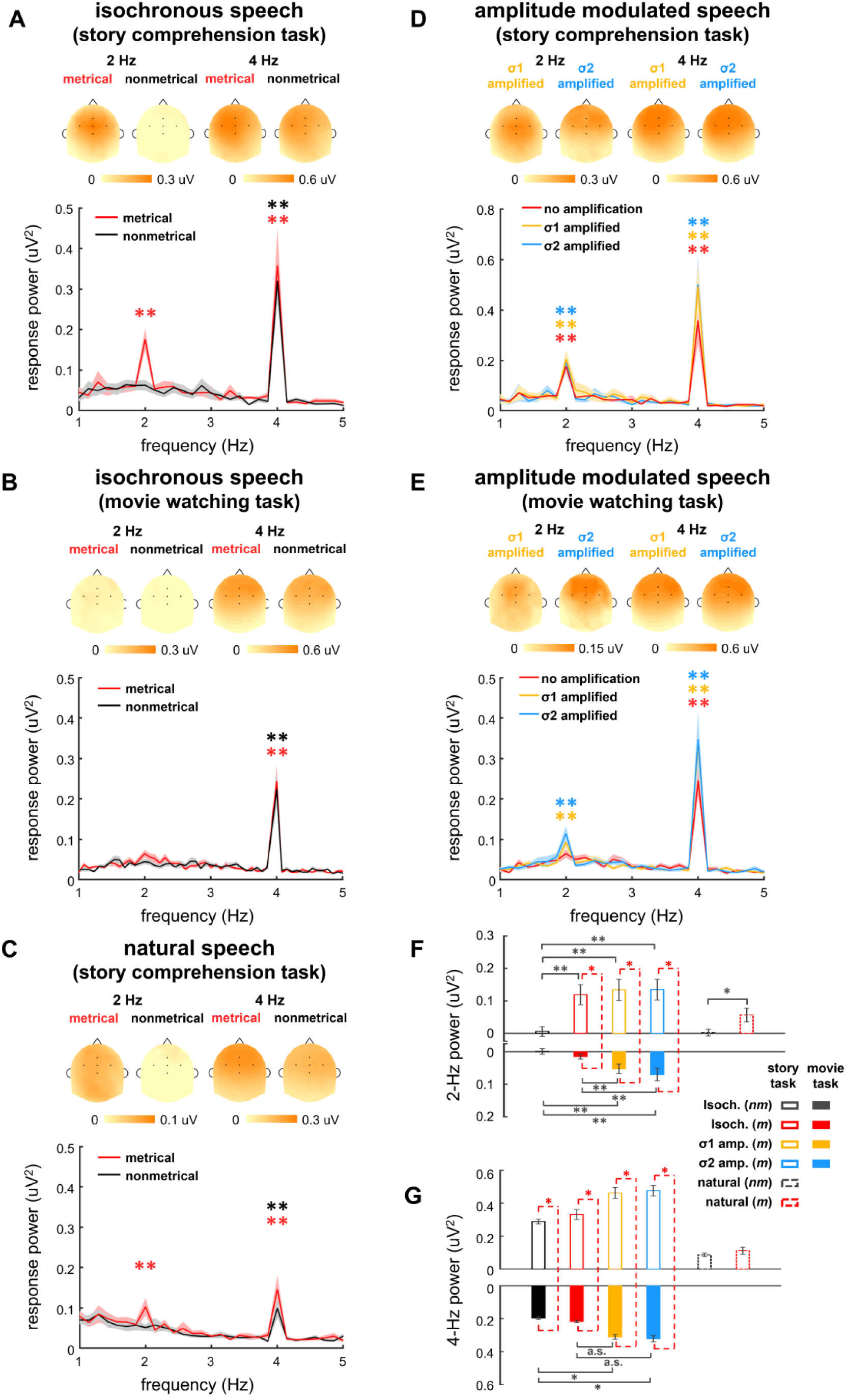
EEG response spectrum. (A-E) Response spectrum averaged over participants and EEG channels. The shaded area indicates 1 standard error of the mean (SEM) across participants. Colored stars indicate frequency bins with stronger power than the power averaged over 4 neighboring frequency bins (2 on each side). The topography on the top of each plot shows the distribution of response power at 2-Hz and 4-Hz. In the topography, the response power is normalized by subtracting the mean power over 4 neighboring frequency bins (2 on each side). The five black dots in the topography show the position of electrode FCz, Fz, Cz, FC3, and FC4. (FG) Normalized power at 2 and 4 Hz. Black stars indicate significant differences between different types of speech stimuli while red stars indicate significant difference between tasks. *P < 0.05, **P < 0.01 (bootstrap, FDR corrected).

Figure 2A showed the responses from participants who attended to stories. For the metrical stories, two peaks were observed in the EEG spectrum, one at 4 Hz, i.e., the syllable rate (P = 0.0001, bootstrap, FDR corrected) and the other at 2 Hz, i.e., the rate of disyllabic words (P = 0.0001, bootstrap, FDR corrected). For the nonmetrical stories, however, a single response peak was observed at the 4-Hz syllable rate (P = 0.0001, bootstrap, FDR corrected), while no significant response peak was observed at 2 Hz (P = 0.36, bootstrap, FDR corrected). When comparing the responses to metrical and nonmetrical stories, a significant difference was observed at 2 Hz (P = 0.0013, bootstrap, FDR corrected) but not 4 Hz (P = 0.32, bootstrap, FDR corrected) (Fig. 2F). The response topography showed a centro-frontal distribution, which was maximal near channel FCz.

When participants watched a silent movie during story listening, however, a single response peak was observed at the syllable rate for both metrical (P = 0.0001, bootstrap, FDR corrected) and nonmetrical stories (P = 0.0001, bootstrap, FDR corrected) (Fig. 2B). The response peak at 2 Hz was not significant for either kind of story (P > 0.056, bootstrap, FDR corrected). When comparing the responses to metrical and nonmetrical stories, there was no significant difference at 2 Hz (P = 0.093, bootstrap, FDR corrected) or 4 Hz (P = 0.29, bootstrap, FDR corrected) (Fig. 2F). These results showed that cortical activity tracked word rhythms during active speech comprehension.

### Neural tracking of words in natural spoken narratives

We next asked whether cortical activity could track disyllabic words in natural speech. The same set of stories used in the isochronous speech condition was naturally read by a human speaker and presented to the participants. The participants correctly answered 95% ± 4% and 97% ± 6% questions for metrical and nonmetrical stories, respectively. In natural speech, syllables were not produced at a constant rate (Fig. 1C), therefore the syllable- and word-tracking response was not frequency tagged. Nevertheless, we could time warp the response to natural speech and make the syllable and word responses periodic. Specifically, the neural response to each syllable in natural speech was extracted and realigned based on a constant 4-Hz rhythm.

The spectrum of the time-warped response was shown in Fig. 2C. For the metrical stories, two peaks were observed in spectrum of the time-warped response, one at the 4 Hz (P = 0.0001, bootstrap, FDR corrected) and the other at the 2 Hz (P = 0.0002, bootstrap, FDR corrected). For the nonmetrical stories, however, a single response peak was observed at the 4 Hz (P = 0.0001, bootstrap, FDR corrected), while no significant response peak was observed at 2 Hz (P = 0.65, bootstrap, FDR corrected). When comparing the responses to metrical and nonmetrical stories, a significant difference was observed at 2 Hz (P = 0.0006, bootstrap, FDR corrected) but not 4 Hz (P = 0.29, bootstrap, FDR corrected). These results demonstrated that cortical activity could track multi-syllabic words during natural speech comprehension.

### Neural responses to amplitude modulated isochronous speech

To investigate potential interactions between neural tracking of words and neural tracking of the speech envelope, we amplitude modulated the isochronous speech at 2 Hz. The amplitude modulation was achieved by amplifying either σ1 or σ2 by a factor of 4 (Fig. 1D), and these two conditions were referred to as the σ1-amplified and σ2-amplified conditions. When listening to amplitude modulated speech, the participants correctly answered 94% ± 12% and 97% ± 8% questions in the σ1-amplified and σ2-amplified conditions, respectively.

The EEG responses to amplitude modulated isochronous speech were shown in Fig. 2DE. When participants attended to speech, a significant 2-Hz spectral peak was observed (σ1-amplified: P = 0.0001, and σ2-amplified: P = 0.0001, bootstrap, FDR corrected) (Fig. 2D). Amplitude modulation did not significantly influence the power of the 2-Hz neural response (σ1-amplified vs. σ2-amplified: P = 0.47; σ1-amplified vs. isochronous: P = 0.35, σ2-amplified vs. isochronous: P = 0.34, bootstrap, FDR corrected) (Fig. 2F). These results showed that the 2-Hz response power was not significantly influenced by 2-Hz amplitude modulation in speech during active speech comprehension.

When participants attended to a silent movie, significant 2-Hz spectral peaks were observed for amplitude modulated speech (σ1-amplified: P = 0.0002, and σ2-amplified: P = 0.0001, bootstrap, FDR corrected) (Fig. 2E). The 2-Hz response was stronger for the amplitude modulated speech than isochronous speech (σ1-amplified vs. isochronous: P = 0.0065, σ2-amplified vs. isochronous: P = 0.0039, bootstrap, FDR corrected) (Fig. 2F), but did not differ between σ1-amplified and σ2-amplified speech (P = 0.23, bootstrap, FDR corrected). Therefore, when attention was diverted, cortical activity tracked the 2-Hz amplitude modulation in speech but not the 2-Hz word rhythm.

### Response phase at 2 Hz

Cortical activity tracking the 2-Hz amplitude modulation was phase-locked to the amplified syllable, and therefore should have a 180° phase lag between the σ1-amplified and σ2-amplified conditions. In contrast, since the σ1-amplified and σ2-amplified conditions had the same word rhythm, the word-tracking response should have the same phase in the two conditions. Therefore, the response phase could provide critical information about whether a 2-Hz response tracked the speech envelope or the word rhythm.

The phase of the 2-Hz EEG response was shown in Fig. 3 for individual participants and individual EEG channels. When participants attended to speech, the 2-Hz response phase was similar between the σ1-amplified and σ2-amplified conditions (Fig. 3). In channel FCz, which showed maximal 2-Hz power (see the 2-Hz topography in Fig. 2), the 2-Hz EEG phase only showed a 19° phase difference between the σ1-amplified and σ2-amplified conditions (95% confidence interval: −45–73°). Furthermore, the phase of the 2-Hz response to isochronous speech was similar to the phase of the response to σ1-amplified (5° phase difference, 95% confidence interval: −52–39°) and σ2-amplified speech (14° difference, 95% confidence interval: −13–39°). Therefore, the 2-Hz EEG response phase was not sensitive to the existence of 2-Hz amplitude modulation in speech or the phase of the 2-Hz amplitude modulation.

**Fig. 3.**
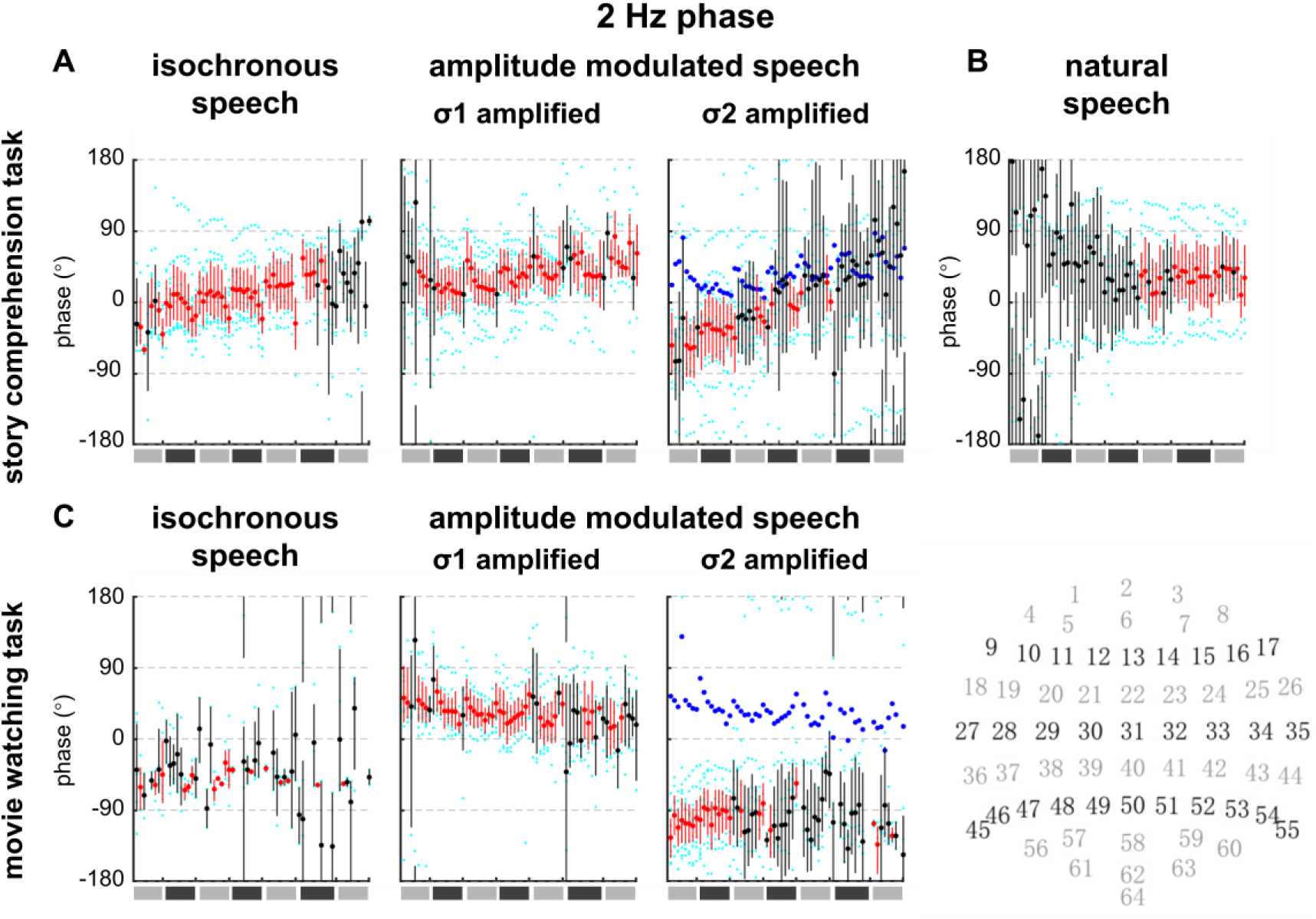
2-Hz response phase in individual EEG channels and individual participants. The x-axis is the EEG channel index, from 1 to 64, which goes from left to right. The approximate scalp position of each channel is shown at the bottom right corner. Individual results are shown as cyan dots. For each EEG channel, the 2-Hz phase averaged over participants is shown by a red or black dot, and the 95% confidence interval across participants is shown by a vertical bar. The dot and bar are red if the inter-participant phase coherence is significantly higher than chance (P < 0.05, see Methods, FDR corrected) and black otherwise. To facilitate the comparison between the σ1-amplified and σ2-amplified conditions, the mean phase in the σ1-amplified condition is repeated in the plot for the σ2-amplified condition as blue dots.

When the participants attended to a silent movie, on the contrary, the 2-Hz response phase showed a clear difference between the σ1-amplified and σ2-amplified conditions. In channel FCz, the 2-Hz phase difference was 139° (95% confidence interval: 98–173°), which was close 180°, consistent with the property of the envelope-tracking response. These results suggested that when attention was diverted, the 2-Hz cortical response predominantly tracked acoustic features of speech.

### Neural tracking of the 2-Hz speech envelope

The response phase analysis demonstrated that the 2-Hz EEG response primarily tracked the word rhythm during speech comprehension but tracked the speech envelope when attention was diverted. This phenomenon, however, could have 2 explanations. One explanation was that the 2-Hz envelope tracking was abolished during speech comprehension. The other explanation was that envelope tracking was enhanced during comprehension but word tracking was more strongly enhanced and therefore dominated the response. To distinguish these explanations, we extracted the neural response component that was phase locked to the 2-Hz envelope (Supplementary Fig. 2A). It was found that there was significant neural tracking of the 2-Hz envelope during both the speech comprehension (P = 0.0008, bootstrap, FDR corrected) and movie watching task (P = 0.0001, bootstrap, FDR corrected), and there was no significant difference between the 2-Hz responses during the two tasks (P = 0.27, bootstrap, FDR corrected).

### Time course of EEG responses to words

The ERP responses evoked by σ1 and σ2 were separately shown in Fig. 4. This analysis was restrained to disyllabic words so that the responses to σ1 and σ2 reflected the responses to the first and second syllables of disyllabic words. When participants attended to speech, the ERP responses to σ1 and σ2 showed significant differences for both isochronous (Fig. 4A) and natural speech (Fig. 4B). When participants watched a silent movie, a smaller difference was also observed between the ERP responses to σ1 and σ2 (Fig. 4A). The topography of the ERP difference wave showed a centro-frontal distribution. For isochronous speech, the ERP latency could not be unambiguously interpreted for isochronous speech since the stimulus was strictly periodic. For natural speech, the ERP responses to σ1 and σ2 differed in a time window between 300-to 500-ms latency. The results for amplitude modulated isochronous speech were shown in Fig. 4CD. When the participants attended to speech, a sharp ERP near the onset of the amplified syllable was observed (Fig. 4C). When the participants attended to the silent movie, the ERP was also stronger for the amplified syllable but the waveform was smoother (Fig. 4D).

**Fig. 4.**
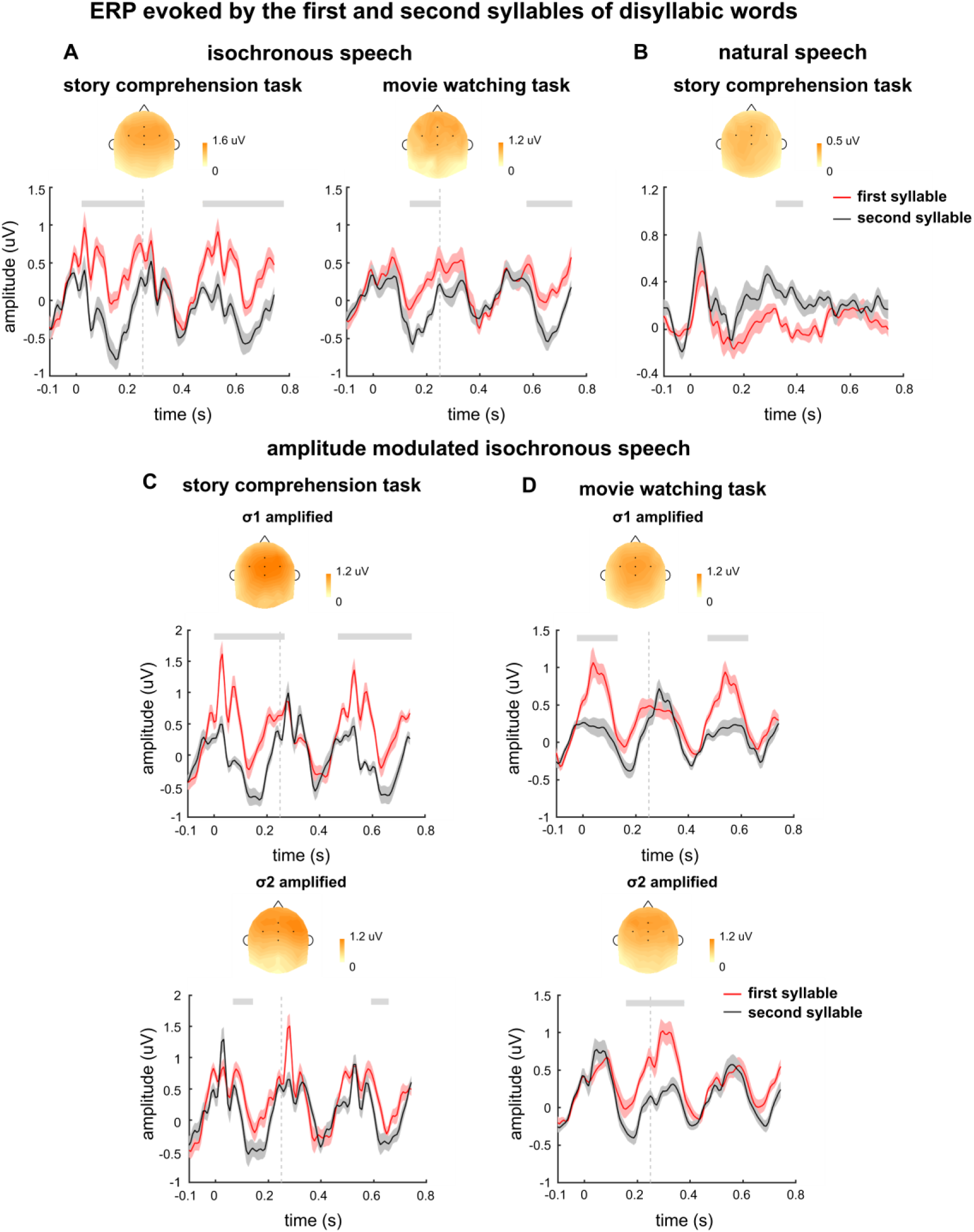
ERP evoked by disyllabic words. The ERPs evoked by the first and second syllable of disyllabic words are shown in red and black respectively. The ERP is averaged over participants and channels. The shaded area indicates 1 SEM across participants. The gray lines on top denote the time intervals in which the two responses are significantly different from each other (P < 0.05, cluster-based permutation test). The topography on top is averaged over all time intervals showing a significant difference between the two responses in each plot. Time 0 indicates the syllable onset.

## Discussion

Speech comprehension is a complex process involving multiple stages, e.g., the encoding of acoustic features and the extraction of linguistic information from acoustic features. Here, we investigate whether low-frequency cortical activity mainly reflects the representation of acoustic features or the representation of linguistic units. It is demonstrated that cortical activity tracks the word rhythm in spoken narratives during a natural speech comprehension task, on top of the speech envelope. Furthermore, using synthesized speech that dissociates the word rhythm and delta-band envelope, it is shown that delta-band cortical activity primarily tracks the word rhythm instead of the envelope during natural speech comprehension. When attention is diverted to a visual distractor, however, delta-band activity becomes synchronized to the speech envelope. These results strongly suggest that linguistic units are critical factors driving large-scale cortical responses to natural speech.

### Neural encoding of linguistic units in natural speech

In speech, linguistic information is organized based on a hierarchy of units, including phonemes, syllables, morphemes/words, phrases, sentences, and discourses. These units span a broad range of time scales, from tens of milliseconds for phonemes to a couple of seconds for sentences and even longer for discourses. How the brain represents this hierarchy of units is a challenging question, and it is an appealing hypothesis that each level of linguistic unit is encoded by cortical activity on the relevant time scale (Ding et al., 2016a; Doumas and Martin, 2016; Giraud and Poeppel, 2012; Goswami, 2019; Keitel et al., 2018; Kiebel et al., 2008; Meyer and Gumbert, 2018).When listening to speech, delta- and theta-band responses are reliably observed (Ding and Simon, 2012; Luo and Poeppel, 2007; Peelle et al., 2013). The time scales of these responses are consistent with the time scales of syllables and larger linguistic units.

Nevertheless, it remains unclear whether these responses directly reflect neural encoding of hierarchical linguistic units or just the acoustic features associated with these units (Daube et al., 2019; Kösem and Van Wassenhove, 2017). On the one hand, neural tracking of sound envelope is reliably observed in the absence of speech comprehension, e.g., when humans listening to unintelligible speech (Howard and Poeppel, 2010; Zoefel and Vanrullen, 2016) and non-speech sound (Lalor et al., 2009; Wang et al., 2012). Neural tracking of the speech envelope can also observed in animal primary auditory cortex (Ding et al., 2016b). On the other hand, evidence for delta-band cortical tracking of linguistic units, such as words and phrases, has been shown using well controlled synthesized speech that removes relevant acoustic cues (Ding et al., 2016a; Jin et al., 2018; Makov et al., 2017). Although synthesized speech can be designed to isolate neural encoding of linguistic units, it remains unclear whether the conclusions from these studies could generalize to natural speech for at least two reasons. One is that, to facilitate frequency-domain analyses, the linguistic units of interests are usually of equal duration, creating a constant rhythm that may be noticed by the listeners. The other is that such synthesized materials usually consists of a list of unrelated linguistic units and therefore the boundaries between units are perceptually more salient than the unit boundaries in natural speech.

The current study demonstrates delta-band cortical activity can indeed track linguistic units, i.e., words, in a coherent spoken narrative, during a natural speech comprehension task, i.e., answering comprehension questions after speech listening. The syllabic- and word-rate responses to natural speech, however, are smaller compared with the responses to isochronous speech, suggesting that strict periodicity in the stimulus can indeed boost rhythmic neural entrainment. Furthermore, the responses to amplitude modulated isochronous speech suggests that, when the word rhythm is in conflict with the speech envelope, cortical activity is primarily phase locked to the word rhythm (Fig. 3). This interpretation also provides a possible explanation for the previous finding that speech comprehension does not always enhance envelope tracking. For example, previous studies have shown that the envelope tracking response is weaker for sentences composed of real words than sentences composed of pseudowords (Mai et al., 2016), and is also for the native language than an unfamiliar language (Zou et al., 2019). It is possible that speech comprehension leads to neural tracking of linguistic units, which competes with and therefore reduces the envelope tracking response.

### Attention modulation of cortical tracking of the speech

It has been shown extensively that cortical tracking of speech is strongly modulated by attention. Most previous studies demonstrate that, in a complex auditory scene consisting of two speakers, attention can selectively enhance the neural tracking of the attended speaker, relative to the neural tracking of the unattended speaker (Golumbic et al., 2013; Horton et al., 2013; Kerlin et al., 2010). These results strongly suggest that the auditory cortex can parse a complex auditory scene into auditory objects, e.g., speakers, and separately represent each auditory object (Shamma et al., 2011; Shinn-Cunningham, 2008). When only one speech stream is presented, cross-modal attention can also modulate neural tracking of the speech envelope but the effect is weaker (Ding et al., 2018; Kong et al., 2014).

Consistent with previous findings, here, the 4-Hz response is also enhanced by cross modal attention (Fig. 2G). The response phase locked to the 2-Hz envelope, however, is not significantly modulated by cross-modal attention (Supplementary Fig. 2A), suggesting that attention does not uniformly enhances all features within the same speech stream. Since the artificially created 2-Hz envelope does not carry linguistic information, the result demonstrates that attention only selectively enhances speech features relevant to speech comprehension. This result extends previous findings by showing that attention can modulate neural tracking of features within a speech stream.

### Time course of ERP response to words

The neurophysiological processes underlying speech perception has been extensively studied using the ERP (Friederici, 2002). Early ERP components such as the N1 mostly reflect auditory encoding of acoustic features, while later components can reflect higher-level lexical, semantic, or syntactic processing (Friederici, 2002, 2012; Sanders and Neville, 2003). In current study, for isochronous speech, response latency cannot be uniquely determined: The syllables are presented at a constant rate of 4 Hz, and therefore a response with latency T cannot be distinguished from responses with latency T ± 250 ms. Furthermore, since the stimulus is periodic, the brain can accurately predict its timing and therefore the responses could be predictive instead of reactive.

For natural speech, the responses to the two syllables in a word differ in a late window with about 400-ms latency. This component is consistent with the latency of the N400 response, which is related to semantic processing of words and can be observed when listening to either individual words or continuous speech (Broderick et al., 2018; Kutas and Federmeier, 2011; Kutas and Hillyard, 1980; Pylkkänen and Marantz, 2003; Pylkkänen et al., 2002). A previous study on the neural responses to naturally spoken sentences has also shown that the initial syllable of an English word elicits a larger N1 and N200-300 than the middle syllable (Sanders and Neville, 2003). A recent study also suggests that the word onset in natural speech elicits a response at ∼100 ms latency (Brodbeck et al., 2018). Language difference is a potential reason why the current study did not observe this early effect: In Chinese, syllables are generally also morphemes while in English most syllables do not carry meaning. The 400-ms latency response observed here is consistent with the hypothesis that the N400 is related to lexical processing (Friederici, 2002; Kutas and Federmeier, 2011). It is also possible that the N400 is weaker for the second syllable since the second syllable in a word is more predictable than the first syllable (Lau et al., 2008).

The difference between the ERPs evoked by the first and second syllables is amplified by attention (Fig. 4AB). Nevertheless, the ERP difference remains significant when participants attend to a silent movie, while the 2-Hz word-rate response in the power spectrum is no longer significant (Fig. 2B). In the response phase analysis, however, it can be observed that the 2-Hz response phase still shows consistency across participants during the movie watching task (Fig. 3). The inter-participant phase consistency is likely to explain why the ERP, which is averaged over participants, can more reliably reflect neural encoding of words. This pattern of results is similar to what is observed in a previous study during unattended listening (Ding et al., 2018), which observes no word-rate response peak in the EEG spectrum but coherent word-rate response phase across participants. These results suggest that word-level processing occurs during the movie watching task, but the word-tracking response is rather weak.

In sum, the current study strongly suggest that cortical activity can track the word rhythm during naturalistic speech listening and delta-band EEG responses are dominated by neural tracking of the word rhythm instead of the speech envelope.

## Materials and methods

### Participants

Forty-eight adults (20-29 years old, mean age, 22.8 years; 27 females) took part in the study. All participants were right-handed native Mandarin speakers, with no self-reported hearing loss or neurological disorders. The experimental procedures were approved by the Research Ethics Committee of the College of Medicine, Zhejiang University (2019-047). All participants provided written informed consent prior to the experiment and were paid.

### Stories

28 short stories were constructed for the study. The stories were unrelated in terms of content, and ranged from 81 to 143 in word count (107 words on average). In 21 stories, the word onset was metrically organized and every other syllable was a word onset. In the other 7 stories, the word onset was not constrained. These two kinds of stories were referred to as metrical stories (Fig. 1A) and nonmetrical stories, respectively (Fig. 1B). Ideally, the metrical stories should be constructed solely with disyllabic words, forming a constant disyllabic word rhythm. Nevertheless, since it was difficult to construct such materials, the stories were constructed with disyllabic words and pairs of monosyllabic words. In other words, whenever a monosyllabic word appeared, it appeared in pairs. After the stories were composed, the word boundaries within stories were further parsed based on a Natural Language Processing (NLP) algorithm (Zhang and Shang, 2019). The parsing result confirmed that every other syllable in the story (referred to as σ1 in Fig. 1A) was the onset of a word. For the other syllables (referred to as σ2 in Fig. 1A), 77% was the second syllable of a disyllabic word while 23% was a monosyllabic word.

In the nonmetrical stories, which were used as a control condition, each sentence also contained an even number of syllables. Nevertheless, no constraints were applied to the word duration, and the odd terms in the syllable sequence were not necessarily a word onset as shown in Fig. 1B.

### Speech

Each story was either synthesized as an isochronous sequence of syllables or naturally read by a human speaker.

### Isochronous speech

All syllables were synthesized independently using the Neospeech synthesizer (http://www.neospeech.com/, the male voice, Liang). The synthesized syllables were 75-354 ms in duration (mean duration 224 ms). All syllables were adjusted to 250 ms by truncation or padding silence at the end, following the procedure in Ding et al. 2016. The last 25 ms of each syllable were smoothed by a cosine window and all syllables were equalized in intensity. In this way, the syllables were presented at a constant rate of 4 Hz (Fig. 1AB). Furthermore, a silence gap lasting 500 ms, i.e., the duration of 2 syllables, was inserted at the position of any punctuation, to facilitate story comprehension.

### Natural speech

The stories were naturally read by a female speaker, who were not aware of the purpose of the study. In the natural speech, syllables were not produced at a constant rate and the boundaries between syllables were labelled by professionals (Fig. 1C). The total duration of speech was 1122 s for the 21 metrical stories and 372 s for the 7 nonmetrical stories.

### Amplitude modulated isochronous speech

Isochronous speech was amplitude modulated to create a 2-Hz speech envelope (Fig. 1D). In the σ1-amplified condition, all σ1 syllables were amplified by a factor of 4. In the σ2-amplified condition, all σ2 syllables were amplified by a factor of 4. Such amplitude modulation was clearly perceivable but did not affect speech intelligibility, since sound intensity is a very weak cue for stress (Zhong et al., 2001) and stress in general does not contribute to word recognition in Mandarin Chinese (Duanmu, 2001).

### Experimental procedures and tasks

The study consisted of three experiments, and each experiment involved 16 participants. Experiment 1 was divided into 2 blocks and within each block the stories were presented in a randomized order. In Experiments 2 and 3, all 28 stories were presented in a randomized order.

#### Experiment 1

The synthesized isochronous speech was presented in Experiment 1. The experiment was divided into 2 blocks. In block 1, the participants listened to isochronous speech including 7 metrical stories and 7 nonmetrical stories. In block 2, participants listened to amplitude modulated speech including 7 σ1-amplified stories and 7 σ2-amplified stories. All 14 stories presented in block 2 were metrical stories and did not overlap with the stories used in block 1. During the experiment, participants were asked to keep their eyes closed while listening to the stories. After listening to each story, the participants were required to answer three comprehension questions by giving oral responses. The person conducting the experiment recorded the responses and then pressed a key to continue the experiment. The next story was presented after an interval randomized between 1 and 2 s (uniform distribution) after the key press. The participants took a break between the two blocks.

#### Experiment 2

The speech stimuli were the same as in Experiment 1, while the task was different. The participants were asked to watch a silent movie (The Little Prince) with subtitles and ignored any sound during the experiment. The stories were presented ∼5 minutes after the movie started, to make sure that participants were already engaged in the movie-watching task. The interval between stories randomized between 1 and 2 second (uniform distribution). The movie was stopped after all 28 stories were presented.

#### Experiment 3

Experiment 3 used the same set of stories as in Experiment 1, but the stories were naturally read. The participants listened to 21 metrical stories and 7 nonmetrical stories. The task was the same as in Experiment 1. The participants took a break every fourteen stories.

### Data recording

Electroencephalogram (EEG) and electrooculogram (EOG) were recorded using a Biosemi ActiveTwo system. Sixty-four EEG electrodes were recorded. Two additional electrodes were placed at the left and right temples to record the horizontal EOG (right minus left), and another two electrodes were placed above and below the right eye to record the vertical EOG (upper minus lower). Two additional electrodes were placed at the left and right mastoids and their average was used as the reference for EEG. The EEG/EOG recordings were low-pass filtered below 400 Hz and sampled at 2048 Hz. The EEG recordings were referenced to the average mastoid recording off-line and band-pass filtered between 0.8 Hz and 30 Hz using a linear-phase finite impulse response (FIR) filter, and the delay caused by the filter was compensated. To remove EOG artifacts in EEG, the horizontal and vertical EOG were regressed out using the least-squares method.

## Data analysis

### Frequency domain analysis

EEG data collected from Experiments 1 and 2 was down-sampled to 128 Hz. Since there was a 500-ms gap between sentences, to avoid the responses to sound onsets, the EEG responses during the 500-ms gaps and to the first two syllables of each sentence were removed from analysis. The EEG responses to the rest of the sentence, i.e., from the third syllable to the last syllable of each sentence, were then concatenated. The concatenated response was 259 s in duration for each condition.

To quantify the consistency of responses across participants, the 259-s EEG responses were divided into 37 trials (∼3 sentences, 7 seconds in duration for each trial). The response averaged over trials was transformed into the frequency domain using the discrete Fourier transform (DFT) without any additional smoothing window. Therefore, the frequency resolution of the DFT analysis was 1/7 Hz. The response power was grand averaged over EEG channels and participants. The inter-trial phase coherence at 2-Hz was separately calculated for each channel and each participant. When averaging the response phase across participants using the circular mean, participants showing no significant inter-trial phase coherence (P > 0.1) were excluded.

Furthermore, the inter-participant phase coherence was calculated for each channel.

### Time domain analysis

This analysis was restrained to the responses to disyllabic words, and the responses to monosyllabic words were not analyzed. The ERPs to the first and second syllables of each disyllabic word were separately extracted and averaged across all disyllabic words. The ERP to each syllable was baseline correlated by subtracting the mean response in a 100-ms window before the syllable onset.

### Time-Warping Analysis

In the natural speech used in Experiment 3, syllables were not produced at a constant rate, and therefore the responses to syllable and word were not frequency tagged. The responses to natural speech, however, could be time warped to simulate the response to isochronous speech. In the time-warping analysis, we first extracted the ERP response to each syllable (from 0 to 750 ms), and simulated the response to 4-Hz isochronous speech using the following convolution procedure:

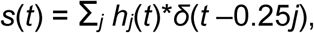

where *s*(*t*) was the time warped response, *δ*(*t*) was the Dirac delta function, and *hj*(*t*) was the ERP evoked by the *jth* syllable, *j* ranging from 1 to the number of syllables in the story. Frequency-domain analysis was then applied to the time warped response, following the same procedure used to analyze the response to isochronous speech.

### Statistical test

In the frequency domain analysis, the statistical tests were performed using bias-corrected and accelerated bootstrap (Efron and Tibshirani, 1994). In the bootstrap procedure, all participants were resampled with replacement 10000 times. To test the significance of the 2-Hz and 4-Hz peaks in the response spectrum (Fig 2A-E), the response amplitude at the peak frequency was compared with the mean power of the neighboring 4 frequency bins (2 bin on each side, one-sided comparison). If the response power at 2-Hz or 4-Hz was stronger than the mean power of the neighboring bins *N* times in the resampled data, the significance level is (*N* + 1)/10001. To test the power difference between conditions within an experiment (solid black lines in Fig. 2F), a two-sided test was used. If the response power was greater in one condition *N* times in the resampled data, the significance level is (2*N* +1)/10001. To test the power difference between experiments (dotted red lines in Fig. 2F), the significance level was *v* if the sample mean in one experiment exceeded the 100-*v*/2 percentile (or fell below the *v*/2 percentile) of the distribution of the sample mean in the other experiment.

To test the phase difference between conditions, the *V*% confidence interval of the phase difference was measured by the smallest angle that could cover *V*% of the 10000 resampled phase difference. In the inter-trial phase coherence test and the inter-participant phase coherence test (Fig. 3), 10000 phase coherence values were generated based on the null distribution, i.e., a uniform distribution. If the actual phase coherence was smaller than *N* of the 10000 phase coherence values generated based on the null distribution, its significance level was (*N* + 1)/10001. When multiple comparisons were performed, the p-value was adjusted using the false discovery rate (FDR) correction (Benjamini and Hochberg, 1995).

In the time domain analysis (Fig. 4), the significant ERP difference between conditions was determined using the cluster-based permutation test (Maris and Oostenveld, 2007). The test was performed following steps given below: (1) The ERP for each participants in two conditions were pooled into the same set. (2) The set was randomly partitioned into two equally sized subsets. (3) At each time point, the responses were compared between the two subsets using a paired t-test. (4) The significantly different data points in the responses were clustered based on temporal adjacency. (5) The cluster-level statistics were calculated by taking the sum over the t-values within each cluster. (6) Steps 2-5 were repeated 2000 times. The p-value was estimated as the proportion of partitions that resulted in a higher cluster-level statistic than the actual two conditions.

### Post-hoc effect size calculation

On top of showing the 2-Hz response power from individual participants and individual channels in supplementary Fig.1, an effect size analysis was applied to validate that the sample size was appropriate to observe the 2-Hz word-tracking response. To simplify the analysis, we calculated the effect size based on a paired t-test to compare the power at 2 Hz and the power averaged over four neighboring frequencies. Since the response power was not subject to a normal distribution, such a t-test had lower power than, e.g., the bootstrap test. However, based on the t-test, the 2-Hz response remained significantly stronger than the mean response averaged over neighboring frequency bins in all conditions shown in Supplementary Table 1. The effect size of the t-test was calculated using the G*Power software (version 3.1) (Faul et al., 2007). We calculated *d* and power based on the mean and standard deviation of the 2-Hz response (reported in Supplementary Table 1). The power was above 0.8 for all conditions, suggesting that the sample size was big enough even for the more conservative t-test.

## Acknowledgements

We thank Dr. Xunyi Pan, Dr. Lang Qin, Jiajie Zou, and Yuhan Lu for thoughtful comments on previous versions of the manuscript. The research is supported by National Natural Science Foundation of China 31771248 (ND), Major Scientific Research Project of Zhejiang Lab 2019KB0AC02 (ND), National Key R & D Program of China 2019YFC0118200 (ND), Zhejiang Provincial Natural Science Foundation of China LY20C090008 (ND), and Fundamental Research Funds for the Central Universities 2020FZZX001-05 (ND).

## Supplementary Figures

**Supplementary Fig. 1.**
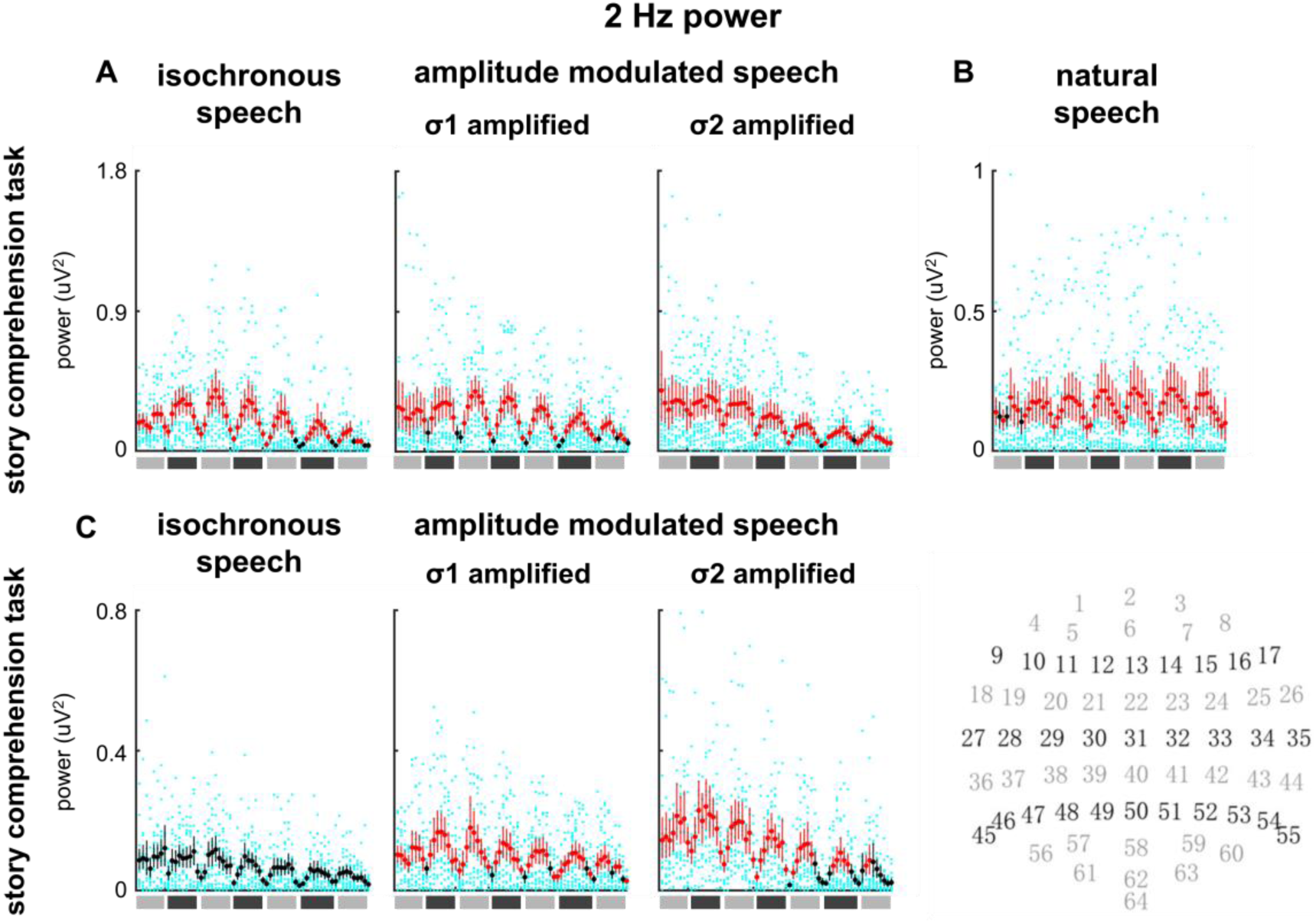
2-Hz response power in individual EEG channels and individual participants. The x-axis is the EEG channels and the channel index, from 1 to 64, which goes from left to right. The approximate scalp position of each channel is shown at the bottom right corner. Individual results are shown as cyan dots. For each EEG channel, the 2-Hz power averaged over participants is shown by a red or black dot, and the 95% confidence interval across participants is shown by a vertical bar. The dot and bar are red if the 2-Hz power is significantly stronger than the power averaged over 4 neighboring frequency bins (P < 0.05, bootstrap, FDR corrected) and black otherwise.

**Supplementary Fig. 2.**
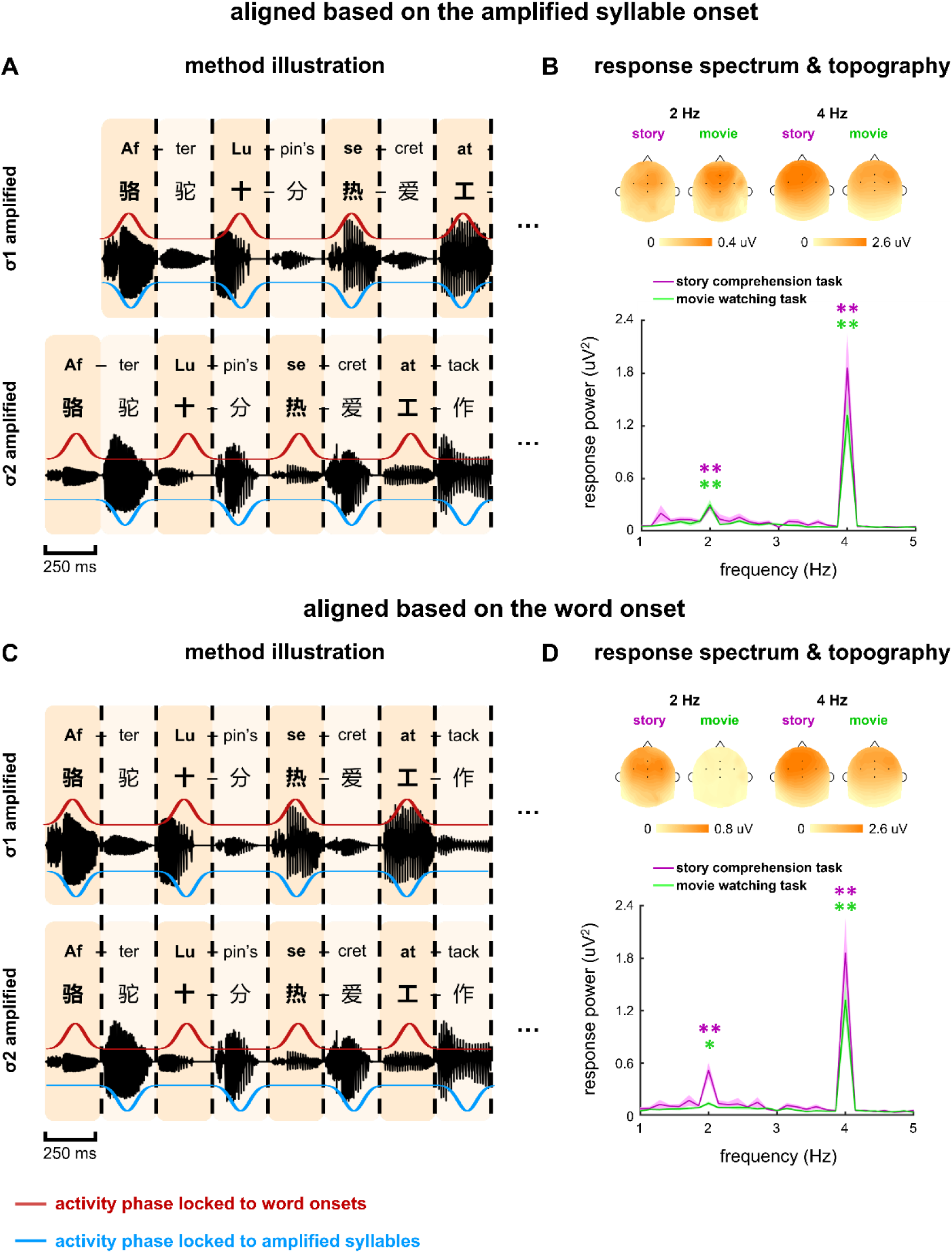
Spectrum of the EEG response aligned to the amplified syllable or word onset. (A) The EEG responses to σ1-amplified and σ2-amplified speech are aligned according to the amplified syllable and averaged. The alignment is achieved by delaying the EEG response to σ1-amplified speech by 250 ms. (B) Spectrum of acoustically aligned EEG response and topography. (C) The EEG responses to σ1-amplified and σ2-amplified speech are directly averaged to extract the response phase locked to the word onset. (D) Spectrum to the word aligned EEG response and topography. *P < 0.05, **P < 0.001 (bootstrap, FDR corrected).

---

**Supplementary Table 1.**

| Condition | *p-value* | *d* | power |
| --- | --- | --- | --- |
| isochronous speech | 0.0016 | 1.21 | 0.99 |
| σ1-amplified speech | 0.0009 | 1.15 | 0.99 |
| σ2-amplified speech | 0.0007 | 1.16 | 0.99 |

**Experiment 2** Effect size of the 2-Hz response for metrical stories
| Condition | *p-value* | *d* | power |
| --- | --- | --- | --- |
| σ1-amplified speech | 0.0026 | 0.97 | 0.95 |
| σ2-amplified speech | 0.0021 | 1.04 | 0.97 |

**Experiment 3** Effect size of the 2-Hz response for metrical stories
| condition | *p-value* | *d* | power |
| --- | --- | --- | --- |
| natural speech | 0.0109 | 0.76 | 0.81 |

